# Smooth Quantile Normalization

**DOI:** 10.1101/085175

**Authors:** Stephanie C Hicks, Kwame Okrah, Joseph N Paulson, John Quackenbush, Rafael A Irizarry, Héctor Corrada Bravo

## Abstract

Between-sample normalization is a critical step in genomic data analysis to remove systematic bias and unwanted technical variation in high-throughput data. Global normalization methods are based on the assumption that observed variability in global properties is due to technical reasons and are unrelated to the biology of interest. For example, some methods correct for differences in sequencing read counts by scaling features to have similar median values across samples, but these fail to reduce other forms of unwanted technical variation. Methods such as quantile normalization transform the statistical distributions across samples to be the same and assume global differences in the distribution are induced by only technical variation. However, it remains unclear how to proceed with normalization if these assumptions are violated, for example if there are global differences in the statistical distributions between biological conditions or groups, and external information, such as negative or control features, is not available. Here we introduce a generalization of quantile normalization, referred to as *smooth quantile normalization* (qsmooth), which is based on the assumption that the statistical distribution of each sample should be the same (or have the same distributional shape) within biological groups or conditions, but allowing that they may differ between groups. We illustrate the advantages of our method on several high-throughput datasets with global differences in distributions corresponding to different biological conditions. We also perform a Monte Carlo simulation study to illustrate the bias-variance tradeoff of qsmooth compared to other global normalization methods. A software implementation is available from https://github.com/stephaniehicks/qsmooth.

## Introduction

Multi-sample normalization methods are an important part of any data analysis pipeline to remove systematic bias and unwanted technical variation, particularly in high-throughput data, where systematic effects can cause perceived differences between samples irrespective of biological variation. Many *global adjustment* normalization methods (Gagnon-Bartsch and Speed 2012; Hicks and Irizarry 2015) have been developed based on the assumption that observed variability in global properties is due to technical reasons and are unrelated to the biology of the system under study (Bolstad et al. 2003; Reimers 2010). Examples of global properties include differences in the total, upper quartile (Bullard et al. 2010) or median gene expression, proportion of differentially expressed genes (Anders and Huber 2010; Robinson and Oshlack 2010; Love, Huber, and Anders 2014), observed variance across expression levels (Durbin et al. 2002) and statistical distribution across samples.

Quantile normalization is a global adjustment normalization method that transforms the statistical distributions across samples to be the same and assumes global differences in the distribution are induced by technical variation (Amaratunga and Cabrera 2001; Bolstad et al. 2003). The observed distributions are forced to be the same to achieve normalization and the average distribution (average of each quantile across samples) is used as the reference.

Several studies have evaluated quantile normalization and other global adjustment normalization methods (Robinson and Oshlack 2010; Bullard et al. 2010; Dillies et al. 2013; Aanes et al. 2014). Under the assumptions of global adjustment normalization methods, quantile normalization has been shown to reduce the variance in observed gene expression data with a tradeoff of inducing a small amount of bias (due to the bias-variance tradeoff) (Bolstad et al. 2003; Qiu, Hu, and Wu 2014). However, when the assumptions of global adjustment normalization methods are violated (for example, if the majority of genes are up-regulated in one biological condition relative to another (Loven et al. 2012; Aanes et al. 2014; Hu et al. 2014; Evans, Hardin, and Stoebel 2016), forcing the distributions to be the same can lead to errors in downstream analyses. Graphical and quantitative assessments (Hicks and Irizarry 2015) have been developed to assess the assumptions of global normalization methods.

If global adjustment methods are found not to be appropriate, another class of normalization methods can be applied (*application-specific* methods), but these often rely on external information such as positive and negative control features or experimentally measured data (Lovén et al. 2012; Aanes et al. 2014). However, it is unclear how to proceed with normalization if the assumptions about the observed variability in global properties are violated, such as they may occur when there are global differences in the statistical distributions between tissues (Figure 1), and external information is not available.

**Figure 1:**
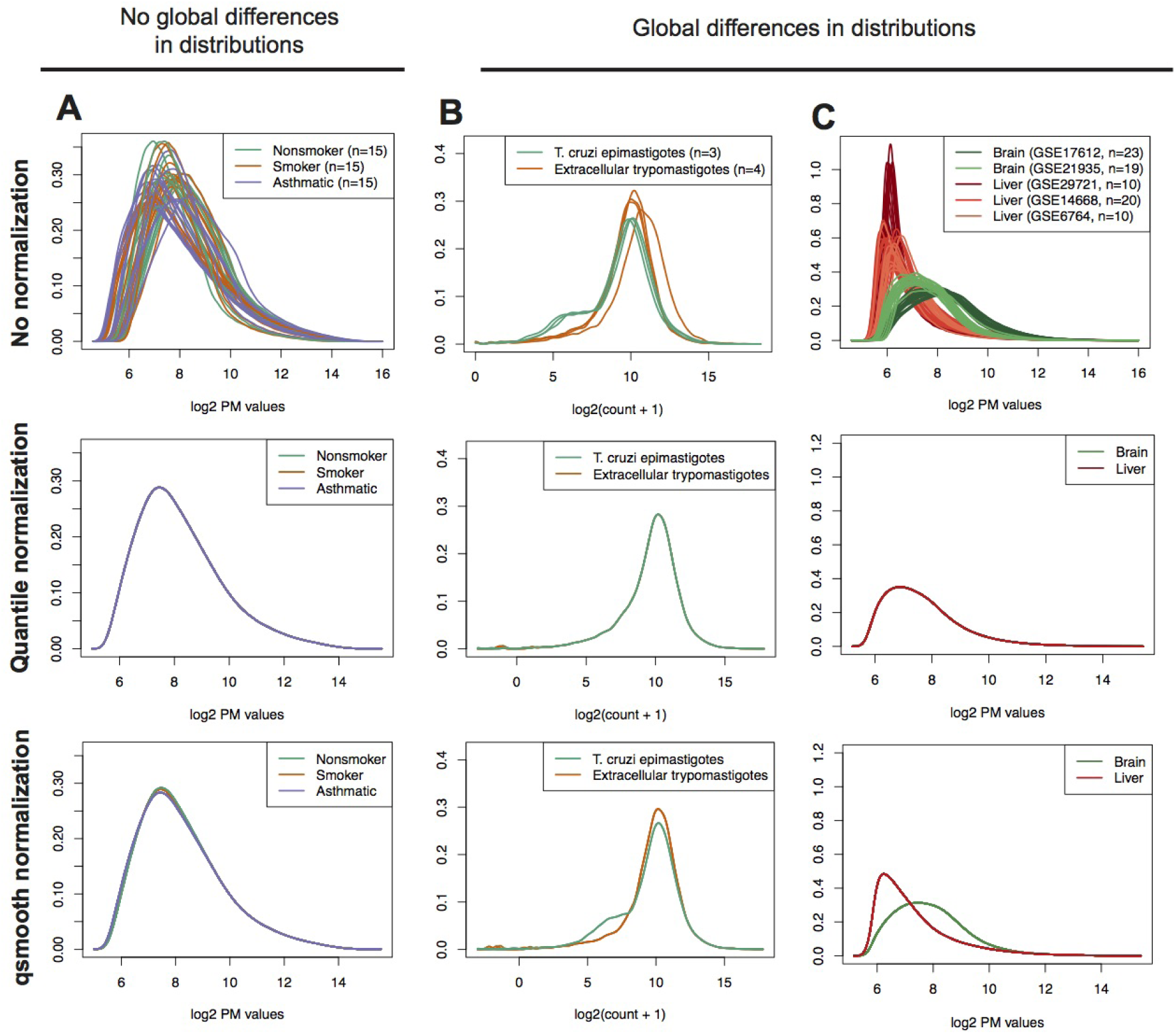
Using biological information to preserve global differences in distributions. Under the conditions of no global differences in distributions (A), qsmooth is similar to standard quantile normalization. Under the conditions of global differences in distributions (B) and (C), quantile normalization removes the global differences by making the distributions the same, but qsmooth preserves global differences in distributions. Examples of gene expression data with (A) Perfect match (PM) values from *n* = 45 arrays comparing the gene expression of alveolar macrophages from nonsmokers (green), smokers (red) and patients with asthma (blue). (B) Gene counts from *n* = 7 from RNA-Seq samples comparing the *T. cruzi* life cycle at the epimastigote (insect vector) stage and extracellular trypomastigotes. Counts have an added pseudocount of 1 and then are log_2_ transformed. (C) PM values from *n* = 82 arrays comparing brain and liver tissue samples colored by tissue (brain [green] and liver [orange]). The shades represent different Gene Expression Omibus (GEO) IDs.

Here we introduce a generalization of quantile normalization, referred to as *smooth quantile normalization* (qsmooth), which is based on the assumption that the statistical distribution of each sample should be the same (or have the same distributional shape) within a biological group (or condition), but that the distribution may differ between groups. At each quantile, a weight is computed comparing the variability between groups relative to the total variability between and within groups (Equation 1). In one extreme with a weight of zero, qsmooth is quantile normalization within each biological group when there are global differences in distributions corresponding to differences in biological groups. As the variability between groups decreases, the weight increases towards one and the quantile is shrunk towards the overall reference quantile (Equation 2) and is equivalent to standard *quantile normalization.* In certain portions of the distributions, the quantiles from different biological groups may be more or less similar to each other depending on the biological variability, which is reflected in the weight varying between 0 and 1 across the quantiles.

Using several high-throughput datasets, we demonstrate the advantages of qsmooth, which include (1) preservation of global differences in distributions corresponding to different biological conditions, (2) non-reliance on external information, (3) applicability to many different high-throughput technologies, and (4) the return of normalized data that can be used for many types of downstream analyses including finding differences in features (genes, CpGs, etc), clustering and dimensionality reduction. We also perform a Monte Carlo simulation study to illustrate the bias-variance tradeoff when using qsmooth.

## Results

### qsmooth: smooth quantile normalization

Consider a set of high-dimensional vectors *Y*_1_, *Y*_2_,…,*Y_n_* each of length *J* representing samples from a high-throughput experiment and each associated with a covariate *Z_i_* representing the biological group or condition. We define 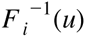 as the empirical quantile function for the *i^th^* sample and the *u^th^* quantile where *u* ∈ [0,1]. Quantile normalization begins by calculating a reference distribution, which is the average at each quantile across the samples, 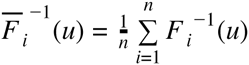. Our method begins by assuming that following form 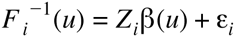. This model is similar to the model described in the functional normalization method proposed by Fortin et al. (Fortin et al. 2014), which relates the quantile functions of a set of high-dimensional vectors to a set of known covariates *Z_i_* that are not associated with biological group or condition. Functional normalization attempts to remove the influence of unwanted technical variation using control features leaving the biological variation in the data. We take a different approach that does not depend on the use of control features and uses a covariate *Z_i_* that is associated with the biological group or condition. In addition, our model extends the model of Fortin et al. by adaptively weighting group information in the normalization transformation applied. Here, 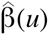 are the estimated regression coefficients representing the reference distributions within each biological group at each quantile and the predicted values, 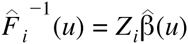 correspond to quantile normalized data within biological groups. We partition the total sum of squares (*SST_(u)_*) into the residual sum of squares (*SSE_(u)_*) and the explained sum of squares (*SSB_(u)_*)

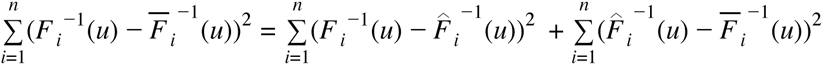

For each quantile *u*, we calculate the weight (*w_(u)_*),

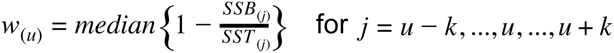

where we use a rolling median across *j* = *u − k,…, u*,…, *u + k* quantiles with a width of *±k* where *k* = *floor*(*N* * 0.05) to smooth the weights at quantiles with a high variance. The number 0.05 is a flexible parameter than can be altered to change the window of the number of quantiles considered. The smooth quantile normalized data is a weighted average,

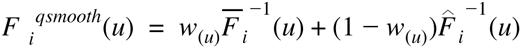

The raw feature values are substituted with the 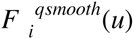 values and then the transformed values are placed in the original order similar to quantile normalization.

### Global differences in distributions between tissues in gene expression and DNA methylation data

We compared qsmooth to other normalization methods using publicly available gene expression and DNA methylation datasets with global differences in distributions. We assessed how global normalization methods impact control features, namely the External RNA Control consortium (ERCC) spike-ins (Jiang et al. 2011), in samples comparing the gene expression from brain and liver tissue in rats (see bodymapRat data set in Methods).

We found that global normalization methods remove the global differences in distribution between brain and liver tissues and induce artificial differences in the spike-in controls compared to using the raw data, including quantile normalization (*p* <2.2e^−16^), Relative Log Expression (RLE) normalization (Anders and Huber 2010) (*p* <2.2e^−16^), and median normalization (*p* <2.2e^−16^) (Figure 2; Supplementary Figures 1–2). In contrast, our method, qsmooth, greatly reduces artificial differences induced between the distributions of the spike-in control genes (*p* = 9.2e^−05^).

**Figure 2:**
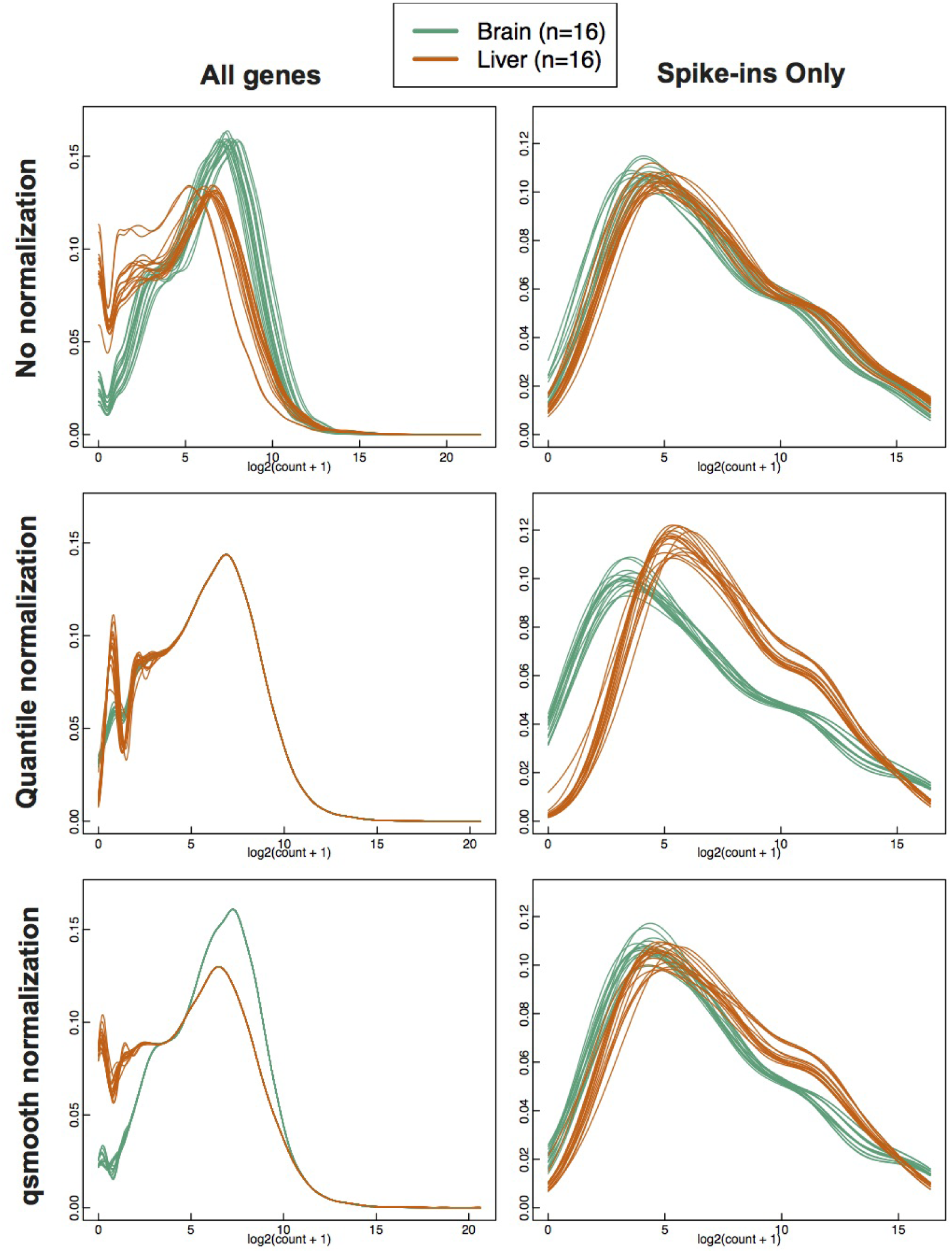
Quantile normalization induces artificial differences in spike-in control genes using data with global differences in distributions. Comparing no normalization (row 1), quantile normalization (row 2) and qsmooth (row 3) applied RNA-Seq gene counts from brain (green) and liver (orange) tissues in the bodymapRat dataset. Column 2 contains the density plots for only the spike-in control genes. Counts have an added pseudocount of 1 and then are log_2_ transformed.

Using the data from the Genotype-Tissue Expression project (GTEx) (GTEx Consortium 2015), we compared qsmooth to a number of scaling normalization methods including, RLE, Trimmed Mean of M-Values (TMM) (Robinson and Oshlack 2010), and upper quartile scaling (Bullard et al. 2010). We observed that scaling methods did not sufficiently control for variability between distributions within tissues; in particular, we observed stark differences in global distribution for a number of body regions, most pronounced between testis, whole blood and other tissues such as artery tibial (Figure 3; Supplemental Figure 3). Normalizing tissues with global differences (in distribution) using a tissue-specific reference distribution, such as in qsmooth, can reduce the root mean squared errors (RMSE) of the overall variability across distributions compared to quantile normalization (Paulson et al. 2016). This occurs because qsmooth is based on the assumption that the statistical distribution of each sample should be similar within a biological group, but not necessarily across biological groups. To demonstrate the importance of preserving tissue-specific differences, we assessed the impact of normalization using quantile normalization and qsmooth using two genes, ENSG00000160882 *(CYP11B1)* and ENSG00000164532 *(TBX20)*. These two genes are known to be highly expressed in specific tissues (Figure 4; Supplementary Table 1). The *CYP11B1* gene has been shown to play a critical role in congenital adrenal hyperplasia (Zachmann, Tassinari, and Prader 1983; Curnow et al. 1993; Joehrer et al. 1997) and the *TBX20* gene plays an important role in cardiac chamber differentiation in adults (Cai et al. 2005; Singh et al. 2005; Stennard et al. 2005; Takeuchi et al. 2005; Qian et al. 2008). In both genes, we found that quantile normalization removes the biologically known tissue-specific expression. In contrast, qsmooth preserves the tissue-specificity, which is also observed just using the raw data. In particular, the *CYP11B1* gene is highly expressed in the testis tissue using both qsmooth normalized and raw data, but it is reported as lowly expressed in the testis tissue after applying quantile normalization. Using qsmooth normalized data and raw data, we observe the tissue-specific gene *TBX20* as highly expressed in heart atrial appendage and heart left ventricle tissues, but lowly expressed in the same tissues after applying quantile normalization. Furthermore, quantile normalization results in this gene being spuriously inflated in other tissues.

**Figure 3:**
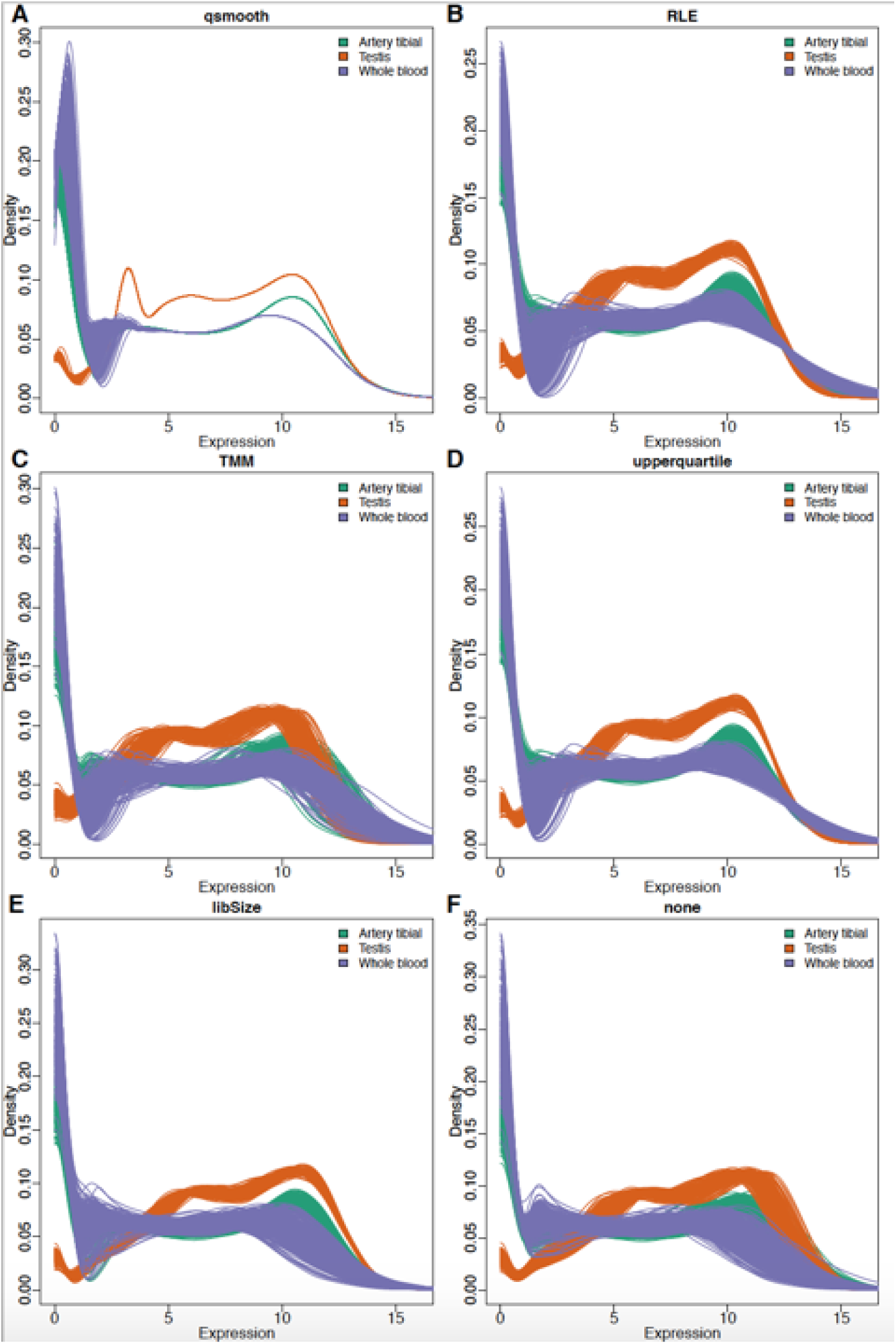
Scaling normalization methods do not adequately control within-group variability. Comparing density plots following either qsmooth (A), Relative Log Expression (RLE) (B), Trimmed Mean of M-Values (TMM) (C), upper quartile scaling (upperquartile) (D), library size (libSize) (E), or no (none) (F) normalization. Plotted are the artery tibial (green) and the testis (orange) tissues from the GTEx consortium. All counts have an added pseudocount of 1 and then are log2 transformed.

**Figure 4:**
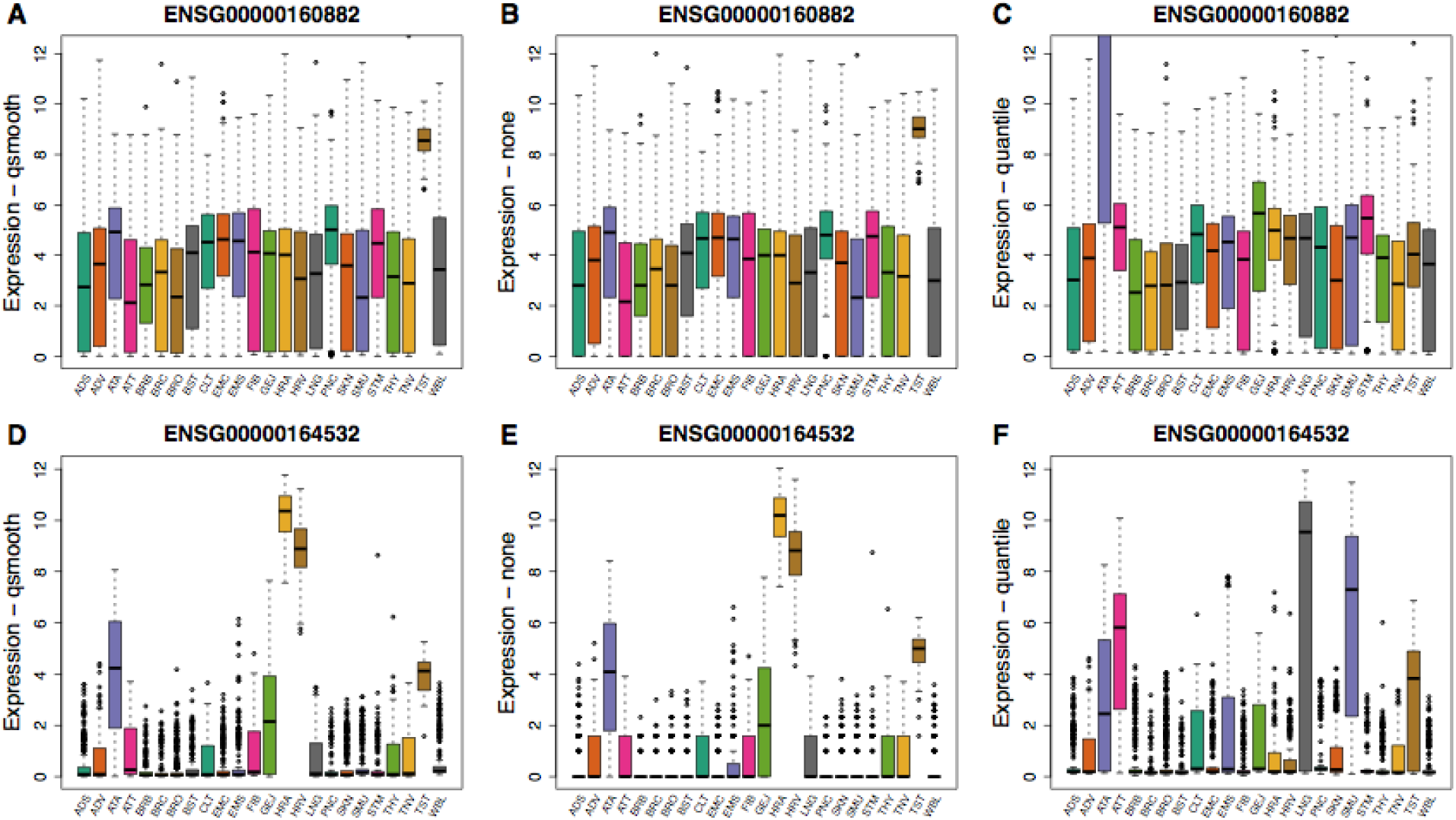
Gene-specific effects induced from quantile normalization. Boxplots of the normalized expression for ENSG00000160882 *(CYP11B1)* and ENSG00000164532 *(TBX20)* are shown for 24 tissues profiled by GTEx. Top, we see *CYP11B1* is more highly expressed in testis (TST) and more lowly expressed in other tissues in both (A) qsmooth and (B) raw expression profiles. However, following quantile normalization (C) *CYP11B1* is relatively lowly expressed in TST but now more variably and highly expressed in the artery aorta (ATA). *CYP11B1* produces 11 beta-hydroxylase, a final step necessary to convert 11-deoxycortisol into cortisol. Steroid 11 beta-hydroxylase deficiency is the second most common cause (5-8%) of congenital adrenal hyperplasia (Zachmann, Tassinari, and Prader 1983; Curnow et al. 1993; Joehrer et al. 1997). Bottom (D, E), *TBX20* is a member of the T-box family and encodes the TBX20 transcription factor and helps dictate cardiac chamber differentiation and in adults regulates integrity, function and adaptation (Cai et al. 2005; Singh et al. 2005; Stennard et al. 2005; Takeuchi et al. 2005; Qian et al. 2008). We see *TBX20* highly expressed in both raw and qsmooth normalized heart atrial appendage and left ventricle tissues (HRA, HRV). However, following (F) quantile normalization, expression of the gene in both heart tissues is almost zero and several other tissues are more highly or variably expressed.

We also tested qsmooth using publicly available DNA methylation (DNAm) data from six purified cell types in whole blood that are known to exhibit global differences in DNAm (Hicks and Irizarry 2015). Using qsmooth, the global differences in distributions are preserved across purified cell types (Figure 5). Furthermore, the cell types cluster more closely along the first two principal components compared to using the raw data or quantile normalized data, because qsmooth accounts for cell type-specific differences in DNAm and removes technical variability across samples within each cell-type.

**Figure 5:**
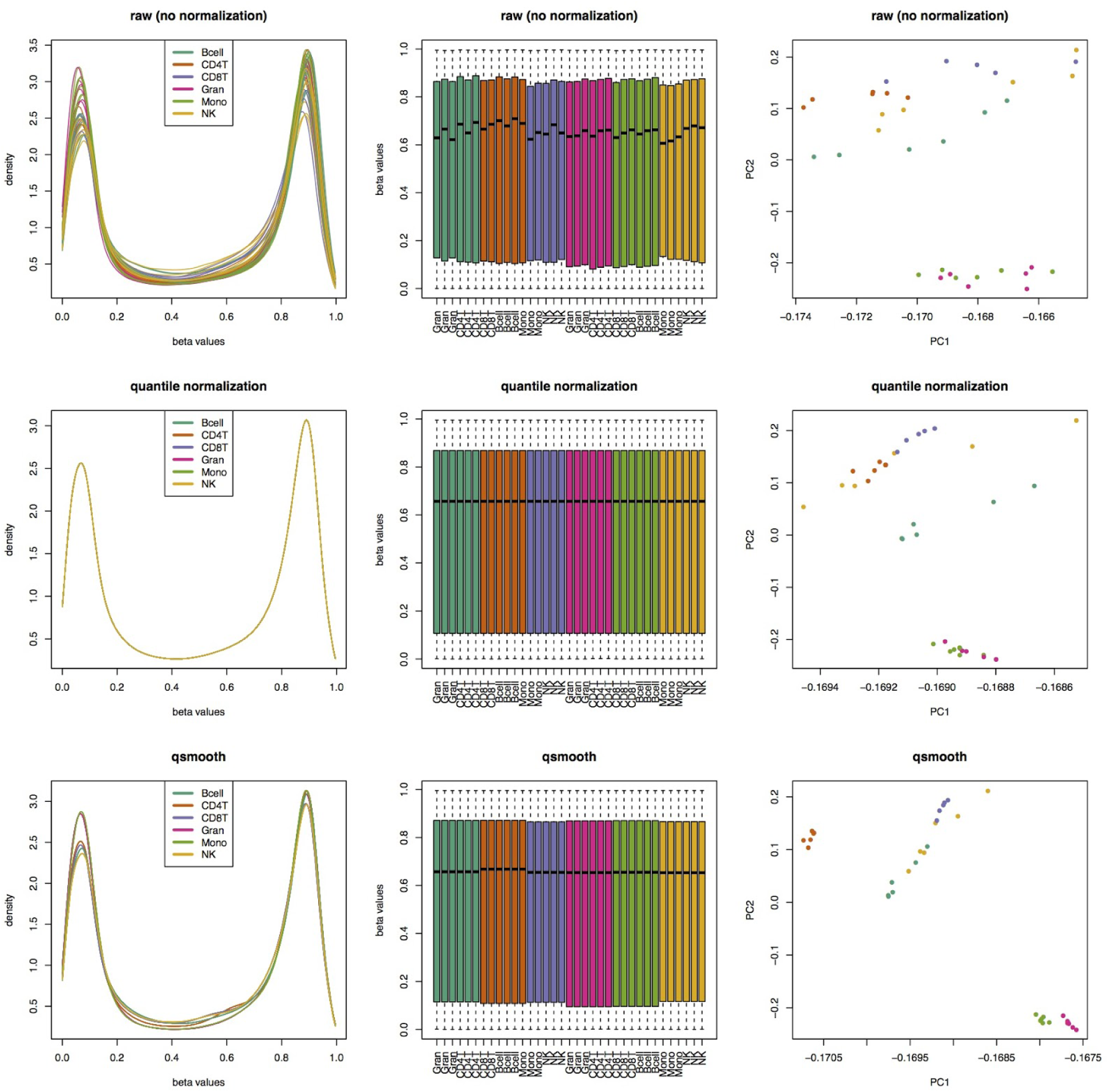
Density plots (column 1) and boxplots (column 2) with global changes in distributions of beta values from n = 35 Illumina 450K DNA methylation arrays comparing raw data (row 1), quantile normalized data (row 2) and qsmooth data (row 3) on six purified cell types from whole blood: CD14+ Monocytes (Mono), CD19+ B-cells (Bcell), CD4+ T-cells (CD4T), CD56+ NK-cells (NK), CD8+ T-cells (CD8T), and Granulocytes (Gran). Column 3 shows first two principal components using three normalization methods.

### The bias-variance tradeoff of qsmooth

We performed a Monte Carlo simulation study to evaluate the performance of qsmooth when the assumptions related to the observed variability in global properties are violated with the detection of differentially expressed genes as a measure of overall performance. We generated gene-level RNA-Seq counts and varied the proportion of differentially expressed genes between biological groups.

As others have noted, when testing for differential expression between groups, quantile normalization results in increased bias with a tradeoff of a reduction in variance compared to using the raw data. Under the assumptions of global normalization methods, qsmooth improves upon this tradeoff, resulting in lower bias compared to quantile normalization, but also less variance compared to using the raw data, and better overall detection of differential expression. As the number of differentially expressed genes increases, quantile normalization and qsmooth both reduce the variance compared to using the raw data, but qsmooth also reduces the bias compared to using the raw data by accounting for global differences between the biological groups, particularly when the assumptions of global normalization methods are violated (Supplementary Figure 4).

## Conclusions

Global normalization methods are useful for removing unwanted technical variation from high-throughput data. However, they are based on the assumption that observed variability in global properties is due only to technical factors and is unrelated to the biology of the system under study. While these assumptions are usually fine when comparing closely related samples, large-scale studies are increasingly generating data where those assumptions do not hold. In cases where these global assumptions are violated, more robust forms of normalization are needed to allow for different distributions in different classes of samples.

Application-specific normalization methods can be applied, but these methods rely on the use of external information such as positive or negative control features or experimentally measured information, which are often not available. Furthermore, these methods are also based on assumptions about the nature of the measured distributions, and these have been shown to be violated in many situations (Dillies et al. 2013; Risso et al. 2014).

The new method we describe here, *smooth quantile normalization* (qsmooth), is based on the assumption that the statistical distribution of each sample should be the same (or have the same distributional shape) within a biological group or condition, but it does not require that different groups or conditions have the same distribution. Our method also does not require any external information other than sample group assignment, it is not specific to one type of high-throughput data, and it returns normalized data that can be used for many types of downstream analyses including finding differences in features (genes, CpGs, etc), clustering and dimensionality reduction.

We demonstrated the advantages of qsmooth using several high-throughput datasets that exhibit global differences in distributions between biological conditions, such as the global changes in gene expression profiles in brain and liver. We illustrated the bias-variance tradeoff when using qsmooth, which preserves global differences in distributions corresponding to different biological conditions. We have implemented our normalization method into the qsmooth R-package, which is available on GitHub (https://github.com/stephaniehicks/qsmooth).

## Methods

### Datasets with global differences in distributions

We downloaded Affymetrix GeneChip gene expression data for alveolar macrophages (GSE2125), brain (GSE17612, GSE21935), and liver (GSE29721, GSE14668, GSE6764) samples in human as reported by a number of studies archived in the Gene Expression Omnibus (GEO) (Edgar, Domrachev, and Lash 2002). We extracted the raw Perfect Match (PM) values from the CEL files using the *affy* (Gautier et al. 2004) R/Bioconductor package for gene expression.

We downloaded raw RNA-Seq gene counts from the *T. cruzi* life cycle (Li et al. 2016). We also downloaded and mapped raw sequencing reads to obtain raw RNA-Seq gene counts for multiple tissues from the Rat BodyMap project (Yu et al. 2014) (GSE53960). This data is also available as an R data package on GitHub, (https://github.com/stephaniehicks/bodvmapRat) (see Supplementary Material for more details). Counts have an added pseudocount of 1 and then are log_2_ transformed. We used the Kolmogorov–Smirnov test for global differences in distributions in spike-ins from the bodymapRat gene expression data.

Gene expression data from the Genotype-Tissue Expression (GTEx) consortium was downloaded from the GTEx portal (http://www.gtexportal.org/) and processed using YARN (Paulson et al. 2016) (bioconductor.org/packages/yarn) (see Supplementary Materials for more details).

The sorted whole blood cell populations measured on Illumina 450K DNA methylation arrays were obtained from *FlowSorted.Blood.450k* R/Bioconductor data package (Jaffe 2015) and the raw beta values were extracted using the *minf* R/Bioconductor package (Aryee et al. 2014).

### Monte Carlo simulation study

We used the polyester R/Bioconductor package (Frazee et al. 2015) to simulate gene-level RNA-Seq counts while varying the proportion of differentially expressed genes (pDiff) to obtain samples with global differences in the distributions between biological conditions. Each simulation study considered ten samples from two groups (total of 20 samples). We added additional non-linear sample-specific noise by splitting the sample into four quartiles and scaling each quartile within the sample with a draw from a uniform distribution ranging from 0.5 to 3. This is more realistic than linearly scaling each sample.

As our measure of performance in the detection of differentially expressed genes, we compared the output of qsmooth to both quantile normalized data and raw (unnormalized) gene counts. We assessed the bias-variance tradeoff of the log_2_ fold change using these three methods while varying the proportion of differentially expressed genes between two groups. The plots were created with the ggplot2 R package (Wickham 2009).

### Software

The R-package *qsmooth* implementing our method is available on GitHub (https://github.com/stephaniehicks/qsmooth).

## Supplementary Materials

Supplementary materials are available in a single pdf, which contain supplemental figures and a detailed description of the bodymapRat and GTEx datasets. All scripts containing the code for these analyses are available on Github (https://github.com/stephaniehicks/qsmoothPaper).

## Acknowledgments

S.C.H and R.A.I. were supported by NIH R01 grants GM083084 and RR021967/GM103552. J.N.P and J.Q. were supported in part by grants 5P01HL105339, 5R01HL111759, 5P01HL114501, 5P50CA127003, 1R35CA197449, 1U01CA190234, 5P30CA006516, and 5R01AI099204. H.C.B and K.O. were supported by NIH R01 grants HG005220 and GM114267.

## Supplementary Material to: Smooth Quantile Normalization

### bodymapRat data

Gene expression data from brain and liver tissues in rat measured using RNA-Seq was obtained from the rat RNA-Seq transcriptomic BodyMap (Yu et al. 2014) (GSE53960), which performed the rat BodyMap across 11 organs and 4 developmental stages. We download the raw FASTQ files and mapped the reads to a custom genome reference made up of the ENSEMBL rat genome (ftp://ftp.ensembl.org/pub/release-80/fasta/rattus_norvegicus/dna), ENSEMBL annotation files (ftp://ftp.ensembl.org/pub/release-80/gtf/rattus_norvegicus/) and the ERCC RNA spike-ins (Jiang et al. 2011) (https://tools.lifetechnologies.com/content/sfs/manuals/ERCC92.zip).

The reads were mapped to the Rat genome using STAR (Dobin et al. 2013) version 2.3.1 with default parameters. Reads mapping to annotated exons were counted using the summarizeOverlaps function in the *GenomicAlignments* R/Bioconductor package. The raw gene counts are available as an ExpressionSet in an R data package on GitHub (https://github.com/stephaniehicks/bodvmapRat). which includes a complete description of processing the raw sequencing reads to obtain gene counts. We filtered out the genes with the sum of counts (across rows) less than the number of samples (columns).

### GTEx data

Gene expression data from the Genotype-Tissue Expression (GTEx) consortium (GTEx Consortium 2015) was downloaded from the GTEx portal website (www.gtexportal.org) and preprocessed using YARN (Paulson et al. 2016). In preprocessing the data we removed samples with very few samples, including the bladder, cervix - ectocervix, cervix - endocervix, fallopian tube, and the leukemia cell line samples. Following the YARN pipeline we used tissues as the biology of the system under study. For most analyses we display only tissues with at least 150 samples. See Supplementary Table 1 for a list of the abbreviations for the tissues.

### Supplemental Tables

**Supplemental Table 1.** Abbreviations for GTEx data

| Tissue | Abbreviation | Subtissue |
| --- | --- | --- |
| Adipose subcutaneous | ADS | Adipose - Subcutaneous |
| Adipose visceral | ADV | Adipose - Visceral (Omentum) |
| Artery aorta | ATA | Artery - Aorta |
| Artery tibial | ATT | Artery - Tibial |
| Brain other | BRO | Brain - Amygdala |
|  |  | Brain - Anterior cingulate cortex (BA24) |
|  |  | Brain - Cortex |
|  |  | Brain - Frontal Cortex (BA9) |
|  |  | Brain - Hippocampus |
|  |  | Brain - Hypothalamus |
|  |  | Brain - Spinal cord (cervical c-1) |
|  |  | Brain - Substantia nigra |
| Brain cerebellum | BRC | Brain - Cerebellar Hemisphere |
|  |  | Brain - Cerebellum |
| Brain basal ganglia | BRB | Brain - Caudate (basal ganglia) |
|  |  | Brain - Nucleus accumbens (basal ganglia) |
|  |  | Brain - Putamen (basal ganglia) |
| Breast | BST | Breast - Mammary Tissue |
| Fibroblast cell line | FIB | Cells - Transformed fibroblasts |
| Colon transverse | CLT | Colon - Transverse |
| Gastroesophageal junction | GEJ | Esophagus - Gastroesophageal Junction |
| Esophagus mucosa | EMC | Esophagus - Mucosa |
| Esophagus muscularis | EMS | Esophagus - Muscularis |
| Heart atrial appendage | HRA | Heart - Atrial Appendage |
| Heart left ventricle | HRV | Heart - Left Ventricle |
| Lung | LNG | Lung |
| Skeletal muscle | SMU | Muscle - Skeletal |
| Tibial nerve | TNV | Nerve - Tibial |
| Pancreas | PNC | Pancreas |
| Skin | SKN | Skin - Not Sun Exposed (Suprapubic) |
|  |  | Skin - Sun Exposed (Lower leg) |
| Stomach | STM | Stomach |
| Testis | TST | Testis |
| Thyroid | THY | Thyroid |
| Whole blood | WBL | Whole Blood |

### Supplemental Figures

**Supplemental Figure 1:**
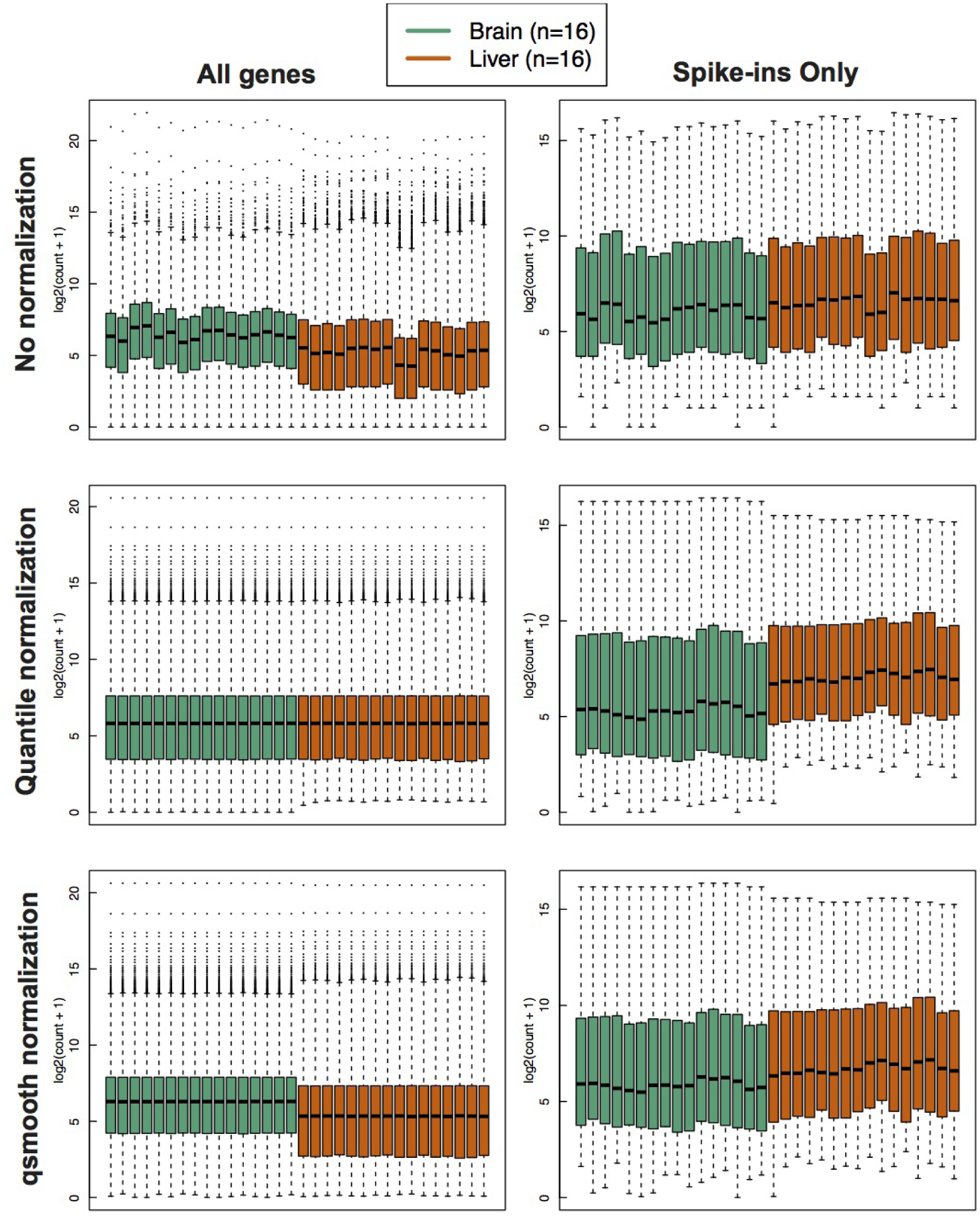
Comparing no normalization (row 1), quantile normalization (row 2) and qsmooth (row 3) applied RNA-Seq gene counts from brain (green) and liver (orange) tissues in the bodymapRat dataset. Column 2 contains the boxplots for only the spike-in control genes. Counts have an added pseudocount of 1 and then are log_2_ transformed.

**Supplemental Figure 2:**
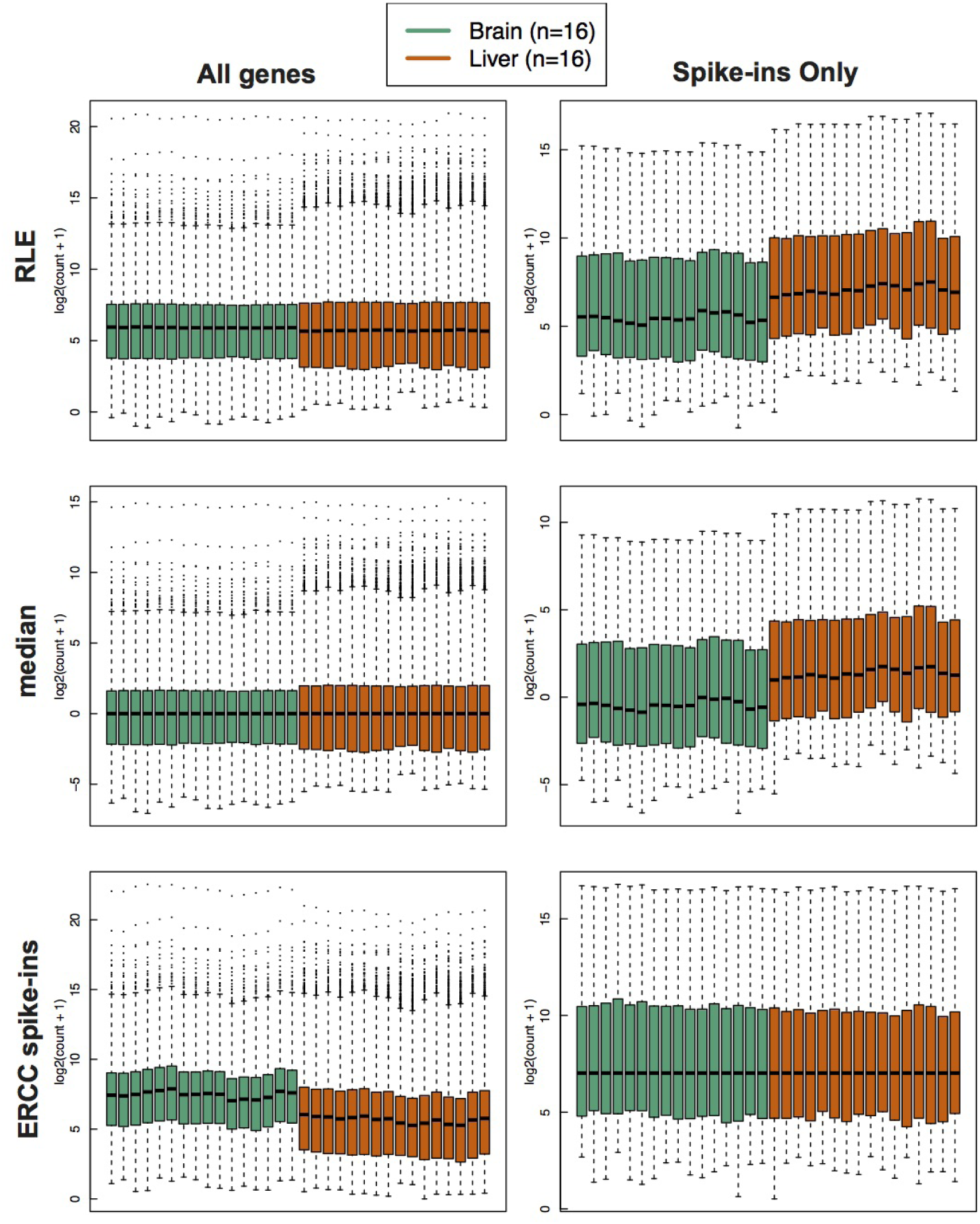
Comparing scaling normalization methods including Relative Log Expression (RLE) normalization (row 1), median normalization (row 2) and scaling by the ERCC spike-ins (row 3) applied RNA-Seq gene counts from brain (green) and liver (orange) tissues in the bodymapRat dataset. Column 2 contains the boxplots for only the spike-in control genes. Counts have an added pseudocount of 1 and then are log_2_ transformed.

**Supplemental Figure 3:**
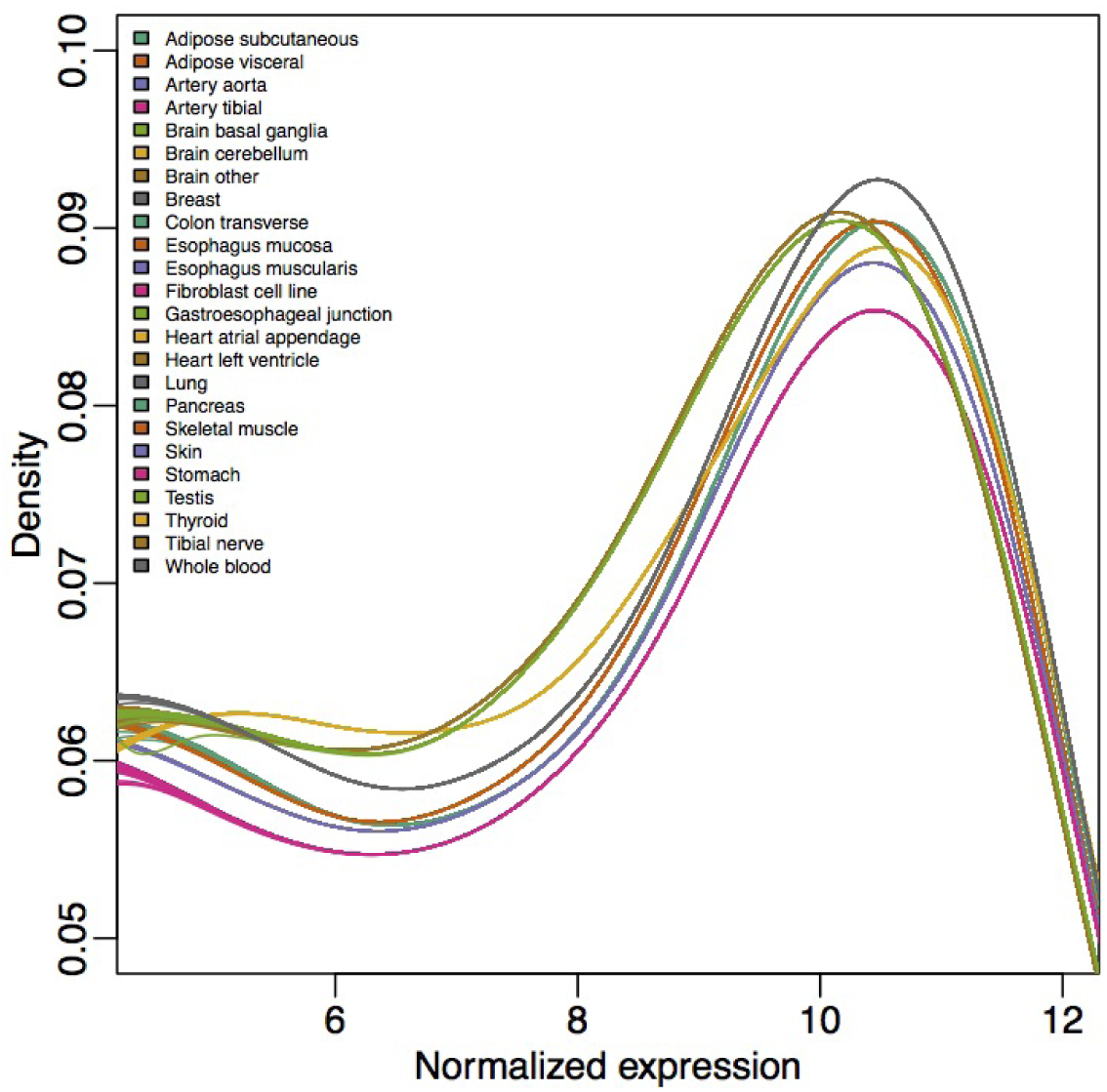
Densities of the gene expression from the GTEx RNA-sequencing samples colored by tissue. Tissue-specific differences in the gene expression distribution can be seen using qsmooth normalization. Only tissues with at least 150 samples are displayed.

**Supplemental Figure 4:**
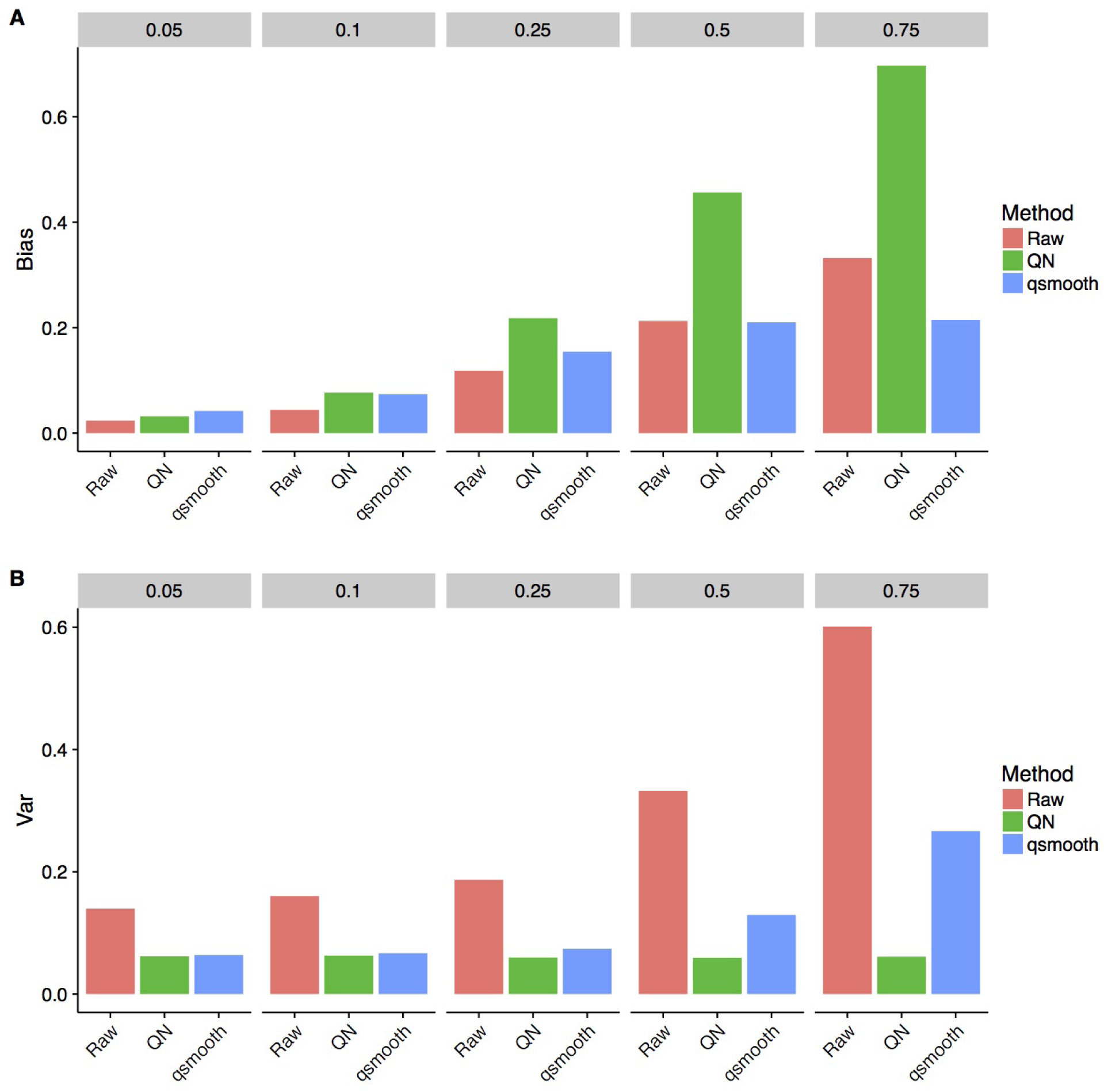
Bias and variance (Var) trade-off of the raw high-throughput data (Raw), quantile normalized data (QN), and smooth quantile normalized data (qsmooth). In each column, we simulated data 10 samples from two biological groups while sampling the proportion of differentially expressed genes (pDiff) from a Uniform[0, X] distribution, where X is listed as the column heading in the figure. Under the assumptions of global normalization methods, qsmooth results in less bias compared to quantile normalization, but also less variance compared to using the raw data. As the number of differentially expressed genes increases, quantile normalization and qsmooth both reduce the variance compared to using the raw data, but qsmooth also reduces the bias compared to using the raw data by accounting for global differences between the biological groups when the assumptions of global normalization methods are violated.

